# Targeted therapy of oncomicrobe *F. nucleatum* with bioengineered probiotic expressing guided antimicrobial peptide (gAMP)

**DOI:** 10.1101/2024.10.29.620919

**Authors:** Ankan Choudhury, Colin Scano, Allison Barton, Christopher M. Kearney, K. Leigh Greathouse

## Abstract

Colorectal cancer (CRC) is a leading cause of cancer-related mortality, with *Fusobacterium nucleatum* (*F. nucleatum*) identified as a key contributor to its progression. This study explores a novel targeted therapy using bioengineered probiotics expressing guided antimicrobial peptides (gAMPs) to selectively inhibit *F. nucleatum*. We engineered *Lactococcus lactis* MG1363 to express gAMPs derived from Ovispirin and Cathelin-related peptide SCF, linked to a Statherin-derived guide peptide (YQPVPE) that binds specifically to the *F. nucleatum* membrane porin FomA.

Our results demonstrate that the Statherin-derived guide peptide enhances the binding affinity to *F. nucleatum*, significantly increasing the preferential attachment compared to control peptides. In vitro assays revealed that both unguided and guided AMPs effectively inhibited biofilm formation in *F. nucleatum*, with gAMPs showing reduced toxicity against non-target bacteria (*Bacteroides fragilis* and *Escherichia coli*). The gAMPs were also more effective in modulating growth kinetics, exhibiting selective toxicity towards *F. nucleatum* at lower concentrations.

Co-culture experiments in a simulated human gut microbiome demonstrated that the gAMP probiotic maintained microbial diversity while effectively reducing *F. nucleatum* abundance. Quantitative PCR and 16S rRNA sequencing confirmed that gAMP treatment preserved the richness of the microbiota, contrasting with significant dysbiosis observed in control samples.

These findings support the potential of engineered probiotics as a targeted therapeutic approach to combat *F. nucleatum*-associated CRC. By leveraging the specificity of Statherin-derived peptides, this strategy not only addresses the pathogenicity of *F. nucleatum* but also mitigates the adverse effects of traditional antimicrobial therapies on beneficial gut microbiota. Future studies will explore the clinical applicability of this approach in CRC management and its impact on overall gut health.

## Introduction

Overall, colorectal cancer (CRC) accounts for approximately 152,810 new cases and 53,010 deaths expected in 2024, making it the fourth most lethal cancer in the U.S. (SEER). While traditionally thought of as a disease of older age, CRC has rapidly become the second leading cause of cancer-related death in people under 50 [1, 2]. The American Cancer Society (ACS) revealed that 20% of cancer diagnoses in 2019 occurred in patients under the age of 55, roughly twice the rate seen in 1995 [3]. Additionally, the incidence of advanced-stage disease in individuals under 50 increased by approximately 3% per year. The ACS projected that in estimated 152,810 new cases and 53,010 deaths are expected in 2024 among those under 50 [3].

Chronic inflammation, diet, and environmental exposures, in combination with somatic mutations, are strong drivers of colon cancer [4]. The microbiome is steadily emerging as a major factor mediating the duration and polarity of the inflammatory response [5]. The oral pathogen, *Fusobacterium nucleatum (F. nucleatum)*, has been identified as a contributor to inflammation, inflammatory bowel diseases, and cancer. Expression of virulence factors, including FomA, FadA, RadD and Fap2 in *F. nucleatum* have established links with invasion of host cells, aggregation, biofilm formation, and trigger multiple pathways of carcinogenesis [6–12]. *F. nucleatum* has consistently been one of the most abundant bacteria in colon adenomas and in colon adenocarcinomas and is associated with colon cancer promotion [13–15]. Moreover, nearly genetically identical strains of *F. nucleatum* are present in both the primary tumor and distant metastases from the same patient with CRC [16]. Recent pioneering research has also uncovered that a particular clade of *F. nucleatum* subspecies *animalis* is the predominant microbe in CRC tumors, which are phenotypically and genotypically distinct from the clades found in the healthy gut microbiota [17]. These clades are also more likely to be highly organized in microniches with immune and epithelial cell functions that promote cancer progression [18]. However, the effects of specifically killing *F. nucleatum* in the presence of a full complement of commensal and pathogenic gastrointestinal microbiota on inflammation, tumorigenesis, and metabolism have been understudied. Therefore, understanding how the removal of *F. nucleatum* from the intestinal environment influences microbial community function is essential for determining its impact on pathogenicity, the early colonization of other oncomicrobes, and the progression of colorectal cancer (CRC) as well as responses to treatment.

Antimicrobial resistance (AMR) is a side effect of cancer treatment, developing in approximately 26% of patients after undergoing chemotherapy and receiving prophylactic antibiotics[19, 20]. Furthermore, AMR is a major issue world-wide, with the most recent data showing approximately 1.27 million deaths attributable to AMR [21–23]. There is an urgent need to overcome this burden by using more targeted approaches like the one proposed in this study, guided antimicrobial peptides (gAMPs or AMPs). Guided antimicrobial peptides (AMPs) hold significant promise in combating multidrug-resistant bacteria, offering a more specific and effective approach to infection control. Unlike broad-spectrum antibiotics, targeted AMPs can be designed to selectively bind to specific microbial membranes, reducing off-target effects and minimizing harm to beneficial bacteria (Xu et al., 2024). This specificity not only enhances efficacy but also mitigates the rise of resistance (Mba and Nweze, 2022). Additionally, AMPs have the advantage of rapid product development compared to the slower pace of traditional antibiotics. The extensive databases available for AMPs and the ease of genetically modifying them for increased specificity, such as through peptide tags, make them an ideal alternative to conventional antibiotics.

One innovative strategy involves using engineered probiotics to express and release guided AMPs (gAMPs). This approach may aid in the recovery of species richness—unlike antibiotic treatments— and reduce overall microbial dysbiosis [24]. *Lactococcus lactis* MG1363, a widely used probiotic, serves as an effective delivery platform and is the first patented engineered probiotic to deliver therapeutic molecules to human patients [25]. Previous research has demonstrated the versatility of *L. lactis* as a delivery system for various antimicrobial peptides, including camel milk lactoferrin derivatives (lactoferrampin and lactoferricin) [26] and gram-negative specific amphibian-derived AMPs [27, 28]. Beyond antimicrobial applications, *L. lactis* has been utilized to deliver systemic therapeutics for conditions such as pulmonary inflammation [29, 30], inflammatory bowel disease [31–34], and more recently, for cancer immunotherapy [35].

Seminal research demonstrates that biofilm formation is a biomarker of colorectal cancer (CRC) and oral squamous cell carcinoma, as well as a potential promoter of local inflammation. *Fusobacterium nucleatum* is a key member of these biofilms. The outer membrane porin, FomA, serves as the anchor binding site on *F. nucleatum* for attachment to the oral mucosa through the human salivary protein statherin. This attachment allows *F. nucleatum* to act as a bridging species for the attachment of secondary colonizers, such as *Porphyromonas gingivalis*, thereby promoting biofilm formation [36]. Targeting FomA with our gAMP offers the additional benefit of reducing biofilm formation at mucosal sites, resulting in a decreased risk of inflammation and disease. Furthermore, this targeting likely exerts added selection pressure to reduce FomA expression in these strains. However, rigorous testing of the downstream corollary effects is essential to ensure no adverse outcomes arise.

To initiate the development of *F. nucleatum*-targeting gAMPs, we utilized the antimicrobial peptide database DBAASP (v3.0) to identify candidate AMPs active against *F. nucleatum*. Ovispirin [37] and Cathelin-related peptide SCF (CPSC5) [38] were selected as promising candidates. Through a comprehensive literature search, we identified a peptide ligand (YQPVPE) derived from the human salivary protein statherin, which specifically binds the *F. nucleatum* membrane porin FomA [39].

We then constructed the gAMPs by linking the ligand to the AMP using a flexible triple glycine linker, resulting in Stat-Ovispirin and Stat-CPSC5 for downstream analysis (Table S1). Synthetic peptides were employed in initial assays to confirm target specific antimicrobial activity. Our results demonstrate that the Statherin-derived guide peptide enhances the binding affinity to *F. nucleatum*, significantly increasing the preferential attachment compared to control peptides. In vitro assays revealed that both unguided and guided AMPs effectively inhibited biofilm formation in *F. nucleatum*, with gAMPs showing reduced toxicity against non-target bacteria (*Bacteroides fragilis* and *Escherichia coli*). The gAMPs were also more effective in modulating growth kinetics, exhibiting selective toxicity towards *F. nucleatum* at lower concentrations.

Building on this foundation, our study focused on engineering *L. lactis* MG1363 to synthesize and deliver gAMPs specifically targeting carcinogenic pathogens. The open-reading frames (ORFs) for both the targeted and non-targeted AMPs were cloned into plasmid pMSP3545 [40], which contains the PNisA promoter [41] inducible by Nisin—an endogenous peptide produced by *L. lactis*. This promoter facilitates controlled expression of the heterologous proteins and peptides. The plasmids were placed downstream of a *L. lactis*-specific secretion signal to create the bioengineered probiotic strains. Nisin-inducible expression systems for heterologous proteins and peptides in both murine and human guts have been well established for over two decades [31, 32, 34, 42, 43], yet their application in delivering guided antimicrobial agents against carcinogenic pathogens remains unexplored. When the resultant bioengineered probiotics were co-cultured with both the target *F. nucleatum*, and other non-target bacteria, the probiotic expressing gAMPs showed targeted toxicity towards *F. nucleatum* with significant reduction in titer of the target bacteria at same probiotic doses compared to the non-target bacteria.

Our approach was inspired by our previous study that developed precision therapy against *Helicobacter pylori* in mice [44]. In that project, *L. lactis* was engineered with plasmids containing open-reading frames (ORFs) for both AMPs and gAMPs—where gAMPs were guided by ligand-peptides targeting the VacA toxin of *H. pylori*. These were expressed under a pH-inducible promoter and secreted in the stomach environment via a secretory signal. The bioengineered probiotic expressing gAMP demonstrated preferential toxicity against *H. pylori* over other tested commensal bacteria in vitro, whereas the probiotic expressing AMP alone did not show this specificity. *In vivo* experiments revealed that both AMP- and gAMP-expressing probiotics effectively reduced *H. pylori* infection and titers in mice when used as either prophylactic or therapeutic agents. Furthermore, microbiome sequencing and analysis indicated that gAMP-expressing probiotics were superior in preserving stomach microbiome diversity compared to their AMP-expressing counterparts.

This study advances the application of engineered *L. lactis* as a targeted delivery system for gAMPs, offering a promising strategy for combating pathogenic bacteria like *F. nucleatum* while maintaining beneficial microbial communities. By specifically targeting biofilm-forming pathogens associated with colorectal cancer and oral squamous cell carcinoma, our approach not only mitigates infection and inflammation but also preserves the integrity of the host microbiome, presenting a significant step forward in precision antimicrobial therapy.

## Materials and Methods

### Bacterial strains and plasmids

The strains of target pathogen *F. nucleatum* use were *F. nucleatum* SB-CTX3Tcol3 derived from CRC tumor biopsy; given to us as a gift from Dr. Suan Bullmann’s group (MD Anderson Cancer Research Center, TX) and *F. nucleatum* subsp. *nucleatum* Knorr (ATCC 25586) derived from Oral lesion; sourced from ATCC (VA). Non-target commensals used for the experiment were *B. fragilis* - *B. fragilis* ATCC 43858 (Enterotoxigenic/ETBF) and *B. fragilis* ATCC 25285 (Non-Enterotoxigenic/NTBF); and *E. coli* K12, both sourced from ATCC (VA). Strains used for synthesis of eGFP and stat-eGFP were *E. coli* 10β and *E. coli* BL21 (DE3) from New England Biolabs (MA). Bioengineered probiotics were created using *L. lactis* subspecies *cremoris* MG1363 (LMBP 3019) acquired from the Belgian Coordinated Collections of Microorganisms (BCCM).

Synthesis of eGFP and stat-eGFP was done using pE-SUMO (Ampicillin) sourced from LifeSensors (PA). The cloning of AMP, gAMP, and sfGFP into probiotic *L. lactis* was done using pMSP3545 sourced from Addgene (MA).

### Synthesis of eGFP and stat-eGFP proteins

The ORFs of eGFP and stat-eGFP were sourced from IDT (CA) as gBlocks, from which they were amplified using PCR and cloned into the pE-SUMO vector using *MfeI* and *BamHI* cut-sites using restriction enzymes and T4 Ligase (Thermo Fisher Scientific). The cloned plasmids were transformed into competent *E. coli* 10β using Heat-shock technique to proliferate the plasmids. Transformed *E. coli*10β were screened by plating on an Ampicillin (50 µg/ml) agar plate and testing the colonies with colony PCR. Positive colonies were propagated in 15 ml Luria-Betani (LB) broth (Ampicillin, 50 µg/ml) at 37°C overnight, pelleted, lysed, and the extracted plasmid was used to transform *E. coli* BL21 (DE3) for protein synthesis. Positively transformed *E. coli* BL21 (DE3) colonies were grown in 2X Yeast-Tryptone (YT) broth in a larger volume (∼500 ml) at 37°C overnight and then induced with 0.5 mM Isopropyl β-d-1-thiogalactopyranoside (IPTG) with a further overnight culture at 16°C. The resultant culture was pelleted at 7000 g and lysed with 0.5 mg/ml Lysozyme along with a freeze-thaw cycle. The thawed lysed pellet is sonicated and centrifuged at 50,000g to obtain the supernatant containing the synthesized proteins, which was purified using Ni-column ion affinity chromatography, utilizing the 6-His tag found in the SUMO fusion partner of the pE-SUMO vector. The purified protein was further cleaved with SUMO Protease, which cleaves the SUMO-fusion partner and releases the synthesized protein.

### Binding assay with Flow Cytometry

Flow cytometry was used to measure cellular binding of the stat-eGFP conjugate protein against the targeted *F. nucleatum* (CRC strain) as well as against the off-target bacteria, *B. fragilis* (ETBF). F*. nucleatum* and *B. fragilis* were both grown anaerobically (5% CO_2_, 5% H_2_, 90% N_2_) in Columbia Broth and Brain-Heart Infusion Broth (BHI) respectively, for 48 hrs or till turbid, at 37°C. The bacteria were then standardized to an OD600 of 1.0 and diluted 1:20 in 1X PBS supplemented with 125 µg/mL unguided eGFP or stat-GFP and incubated for 30 minutes at 37°C, shaking at 180 rpm. Cells were washed and resuspended in 1× PBS and flowed through a BD FACSVerse system (BD Biosciences, NJ). Cells were excited with a blue 488 nm laser using a 488/10 bandpass filter. For each sample, fluorescence intensity measures were recorded for a total of ≥4,000 events collected in triplicate. Data were analyzed using FCS Express (De Novo Software, CA). Statistical analyses were done for both the median fluorescence and percent of events positive for eGFP by using ANOVA.

### Biofilm Assay with synthetic AMP/gAMPs

Synthetic peptides (HPLC purified) for CPSC5, Ovispirin and their Stat-guided analogs were purchased from ABI Scientific (VA). After reconstitution in sterile PBS buffer, the peptides were added to a 96 well plate in a concentration gradient from 0.5 to 32 µM in appropriate media for the bacteria of choice (Columbia for *F. nucleatum*, BHI for *B. fragilis*, LB for *E. coli*) up to a volume of 0.2 ml in each well. The bacteria, grown in appropriate media either anaerobically or aerobically, was standardized to OD600 of 1.0 and 5 µl of it were added to each well, followed by incubation for appropriate amount of time (overnight aerobically for *E. coli*, 72 hr anaerobically for *F. nucleatum*, 48 hr anaerobically for *B. fragilis*). The plate post-incubation was assayed for biofilm formation using a modifed protocol described by O’ Toole et al [45]. Briefly, the media was aspirated from the wells by pipetting, followed by fixing the biofilm with 0.2 ml of Methanol added to each well, evaporation of the methanol by aeration/incubation at 50°C, addition of equal volume of 1% Crystal Violet solution, 3X washing by water or till the wash runs clear, dissolution of the crystal violet bound to the biofilm in 0.2 ml of 30% Acetic acid solution, transfer of the crystal violet-acetic acid solution to a fresh 96 well plate for colorimetric assay at 600 nm. One row of the plate was left only to grow the bacteria without any AMP/gAMP added, the absorbance reading of that row is taken as the positive control, and biofilm inhibition % is calculated for every other well from that control.

### Growth kinetics assay with synthetic AMP/gAMPs

For growth kinetics assay, the AMP/gAMPs and the bacteria were set up in a 96 well plate in the identical manner as the biofilm assay. The incubation in this case was done inside the Stratus Kinetic Microplate Reader (Cerillo, VA) at 600 nm to read the cell growth kinetics. The data post-incubation period was procured and analyzed using the Cerillo Labrador Software, with urther statistical representation done in R.

### Cloning antimicrobial peptides and guided antimicrobial peptides in *L. lactis*

The ORFs of the AMPs (**Table S1**), codon-optimized for *L. lactis*, were cloned into the pMSP3545 plasmid (**Fig 4a**) which plasmid includes the PNisA Nisin-inducible promoter and the usp45 signal peptide for extracellular secretion of the downstream peptide. The ORFs were amplified from a gBlock by PCR (1 minute melting at 95°C; 35 cycles of 15 seconds melting at 95°C, 15 seconds annealing at Tm + 3°C, 30 seconds extension at 72°C; and 5 minutes elongation at 72°C) using respective primers and pasted into the pMSP3545 plasmid using restriction enzyme cut sites post agarose gel purification. The recombinant plasmid with AMPs/gAMPs were electroporated into electrocompetent *L. lactis* MG1363 (LMBP 3019) cells. Briefly, the procedure involved mixing of thawed electrocompetent *L.lactis* with at least 0.25 µg of plasmid, resuspended in ice-cold 0.5 M sucrose and 10% glycerol, electroporation in a cuvette (2000 V, 25 µF, 200 Ω) for a pulse of 5 ms, and grown out at 30°C in GM17 media (M17 media with 0.5% Glucose). The screening for positive transformation was done by plating on GM17 agar plate with erythromycin (5 µg/ml).

### Co-culture assay of probiotic with bacteria monoculture

*L. lactis* clones engineered to express AMP or gAMP were propagated from glycerol stocks and grown in GM17 broth overnight at 30°C with erythromycin (5 µg/ml) with no shaking. *F. nucleatum* was grown in Columbia broth anaerobically at 37°C for 48-72 hrs. The *L. lactis* cultures were serially diluted in a 96-well culture plate with GM17 broth to make up a volume of 100 µL. To each well, 10 µL of the *F. nucleatum* culture were added, and each well volume was brought up to 200 µL with Columbia broth. The plate was left to grow overnight in anaerobic conditions. After 24 hours, the well contents were transferred to a 96-well PCR plate. That PCR plate was sealed, heated for 15 minutes at 100°C, and then chilled at 4°C for 5 minutes. This plate was then centrifuged at 2,000 *g* for 2 minutes, and the supernatant was used as the template for qPCR.

qPCR was carried out using primers for the FomA gene (forward: 5’-ACTTTACCAGTTGCCCAGTT-3’; reverse: 5’-GGAGACCAAATGGTTCAGTAGAT-3’)to quantify *F. nucleatum* titer and primers flanking the *L. lactis acma* gene (forward: 5′-GGAGCTCGTGAAAGCTGACT-3′; reverse: 5′-GCCGGAACATTGACAACCAC-3′) were used to quantify *L. lactis* titer. The qPCR used SYBR Green as the amplification dye and ROX as the passive dye, and the thermal cycler had 2 minutes melting at 95°C followed by 40 cycles of 15 seconds melting at 95°C, and 1 minute of annealing/extension at 60°C, ending with a melt curve. Standard curves for *F. nucleatum* and *L. lactis* were constructed by determining C_T_ values from the qPCR data for different dilutions of the gDNA extracted from the cultures of the respective bacteria (1/10, 1/100, 1/10^3^, and 1/10^4^) in the qPCR plates, and the approximate number of genomes represented in the gDNA was calculated by dividing the amount of gDNA by the average size of the genome of that species. The same procedure was followed with the off-target bacteria where *B. fragilis* and *E. coli* were co-cultured with serially diluted cultures of *L. lactis* for 24 hours, and the titers of the off-target bacteria were determined by qPCR using primers for species-specific genes for either bacterium, DE3-T7 polymerase for *E. coli* (forward: 5’-GAAGCTTGCTTCTTTGCT-3’; reverse: 5’-GAGCCCGGGGATTTCACAR-3’)and *Bft*/Bf toxin for *B. fragilis* (forward: 5’-GGTTTCAACCGTCAGGTACA-3’; reverse: 5’-GCGAACTCATCTCCCAGTATAAA-3’). The amount of *L. lactis* added to the co-cultures of all the three assays ranged from approximately 2.8×10^6^ to 1.8×10^8^ CFU/ml. The statistical difference between any data sets was performed by ANOVA.

### Co-culture assay of probiotics with polymicrobial community culture

Three human fecal samples were randomly selected from the control arm of our vitamin D microbiome randomized control trial (The study was approved by the Institutional Review Board (IRB) (1845028-2) at Baylor University and registered under the identifier NCT05387876 at ClinicalTrials.gov.) The polymicrobial community was created by reanimating fecal samples from human subjects kept frozen at −80°C in the complex media described by Li *et. al*. for *ex vivo* microbiome culture in their MiPro model [21], in anaerobic condition at 37°C for 48 hrs. These polymicrobial cultures were added in 96 well (2 ml, deep well) with *F. nucleatum* and probiotic of choice in the ratio 6:1:1. For samples that were *F. nucleatum* only negative control, the rest of the volume was made up by adding more of the complex media. The plates were incubated anaerobically at 37°C with separate plates that served as 0 hr, 24 hr and 48 hr timepoint samples. Post incubation, the samples were extracted using Quick DNA Fecal/Soil Microbe 96 Kit (Zymo Research Corp, CA). The extracted gDNA was either subjected to 16s rRNA Illuminq sequencing or *F. nucleatum* specific sequencing as described in the previous protocol.

### Illumina 16S rRNA sequencing of polymicrobial samples

The extracted gDNA was sequenced as per the protocol described in the Earth Microbiome Project [46]. The 16S rRNA variable region V4 was amplified with polymerase chain reaction (PCR), using the following primers, 16S Forward Primer (515 F) with adapters: TCG TCG GCA GCG TCA GAT GTG TAT AAG AGA CAG GTGYCAGCMGCCGCGGTAA 3’, and 16S Reverse Primer (926 R) with adapters: 5’ GTC TCG TGG GCT CGG AGA TGT GTA TAA GAG ACAG CCGYCAATTYMTTTRAGTTT 3’.The PCR mixture comprised 1 μl of each forward and reverse primer (12.5 μM), 5 μl of extracted DNA of approximately equal concentration from each sample, 12.5 μl of 2x Platinum Hot Smart Master Mix (Thermo Fisher Scientific), and water to make a final volume of 25 μl. The amplifications were performed under the following conditions: initial denaturation at 94 °C for 5 min, followed by 35 cycles of denaturation at 94 °C for 45 s, primer annealing at 50 °C for 1 min, and extension at 72 °C for 1 min 30 s, with a final elongation at 72 °C for 5 min. The presence of PCR products was visualized by electrophoresis using a 1.5% agarose gel. The products were cleaned using Sera-Mag select PCR clean up kit (Cytiva). A second PCR was conducted to attach the Illumina adapters and barcode index primers-I5 (forward) and I7 (reverse). After adapter and index attachment, the amplicons were normalized and pooled together in a DNA library at a concentration of 4 nM, measured by Quant-iT dsDNA Assay Kit (Thermo Scientific). The pooled DNA library was paired end sequenced at 2×300 bp using a MiSeq Reagent Kit v3 on Illumina MiSeq platform (Illumina, CA), at Baylor University.

### 16S rRNA sequencing data analysis

The sequences were evaluated, demultiplexed, and filtered using QIIME2 (version 2024.5) [47] plugins on the Kodiak High-Performance Computing (HPC) Cluster hosted by High Performance and Research Computing Services, Baylor University. Both paired reads were trimmed from the forward end and read length of at least 200 bp for further processing to generate the ASVs. The denoising and filtering of chimeric sequences was done using the DADA2 plug in of QIIME 2. The samples were rarefied at a depth of >2500 which retained 97% of the samples. Taxonomic classification was performed utilizing the QIIME2-compatible pre-trained feature classifier based on 16S rRNA gene database from Silva 16S rRNA 138 database. The alpha diversity (Shannon, Faith Phylogenetic Diversity, and Species Richness) and beta diversity (Bray-Curtis, Aitchison’s, Weighted and Unweighted Unifrac) analyses were performed using QIIME2 plugins. The resultant α-diversity, β-diversity, and ASV tables were downloaded and further analyzed in R 4.3.1 and RStudio platform 2023.12.1. The resultant taxonomic abundance data (at genus level) was analyzed using the CCREPE package in R [48] to determine the correlation between the taxa according to relative changes in abundance upon addition of *F. nucleatum*. The resultant coterie of genera (|correlation|>0.25, P-value < 0.05, FDR adjusted P-value <0.15, iterations = 1,000) was then used to construct the equation for Dysbiosis Index Equation. Every other data visualization used default tools in the *tidyverse* and/or *ggplot2* packages.

## Results

### The Statherin derived guide peptide preferentially attaches to *F. nucleatum*

The goal of our initial experiments was to determine if our guide peptide had specificity towards FomA on *F. nucleatum.* The binding affinity of the statherin-derived guide peptide ligand was demonstrated by synthesizing eGFP fused with the peptide ligand (stat-eGFP) and quantifying the binding with flow cytometry post-incubation with *F. nucleatum* and *B. fragilis* (non-*Fusobacterium* candidate). Both the median fluorescence and the percent of fluorescent positive events were significantly higher (p<0.01, ANOVA) for *F. nucleatum* incubated with stat-eGFP than with either native eGFP or untreated bacteria (**Fig. 2A-2B**). The fluorescence values from the same bacteria treated with native eGFP was higher than the untreated sample but not significantly. In contrast, *B. fragilis* treated with either native or stat-guided eGFP exhibited no significant difference between themselves (p=0.58, ANOVA) although they were more significantly fluorescence than the untreated samples (p<0.05, ANOVA). Thus, we show here that eGFP with the stat-guide ligand peptide preferentially attaches to the *F. nucleatum* cells compared to native eGFP, unlike in non-target bacterium *B. fragilis*. Our data indicate that the ligand derived from the statherin (stat) peptide is an effective guide for fluorescent molecules to attach to the target pathogen *F. nucleatum* preferentially over unguided molecules, which was hindered in commensal anaerobes.

**Fig 1.**
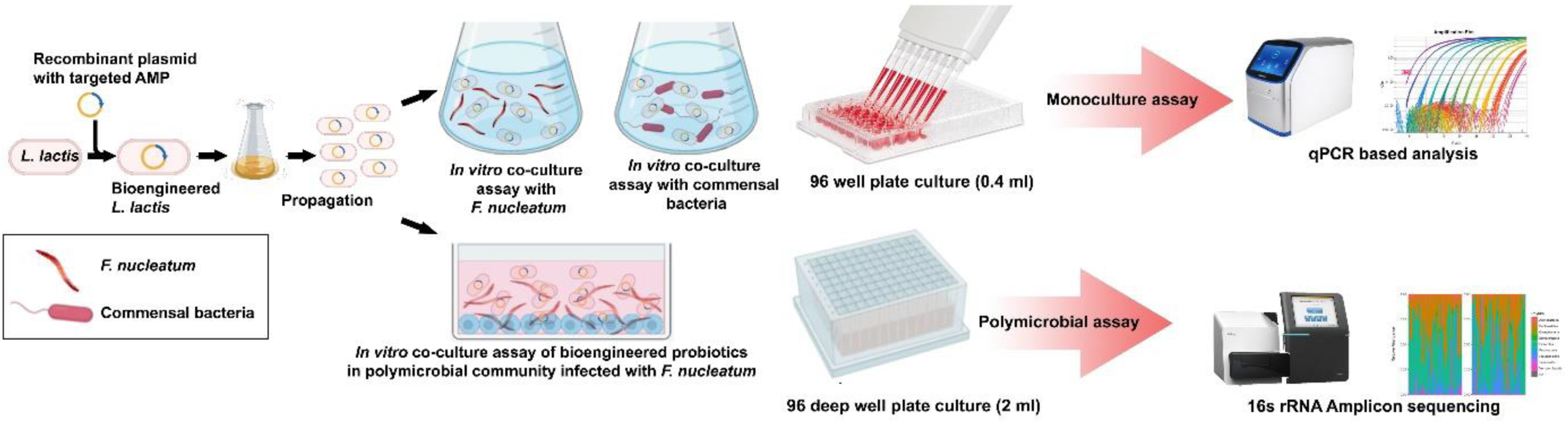
Experimental Overview. The probiotic *L. lactis* was cloned with a Nisin-inducible plasmid with the ORF of the AMP/gAMP downstream of it, to express and secrete the gAMP in vitro where the statherin derived guide peptide will allow the AMP to preferably attach to the FomA membrane protein on *F. nucleatum* and exhibit targeted antimicrobial activity. The bioengineered probiotic was assayed by either co-culturing with a single *Fusobacterium* and non-*Fusobacterium* species and analyzing by species specific qPCR (top) or co-culturing with a fecal microbiome derived polymicrobial community spiked with *F. nucleatum* and analyzing through 16s rRNA metagenomics (bottom).

**Fig 2.**
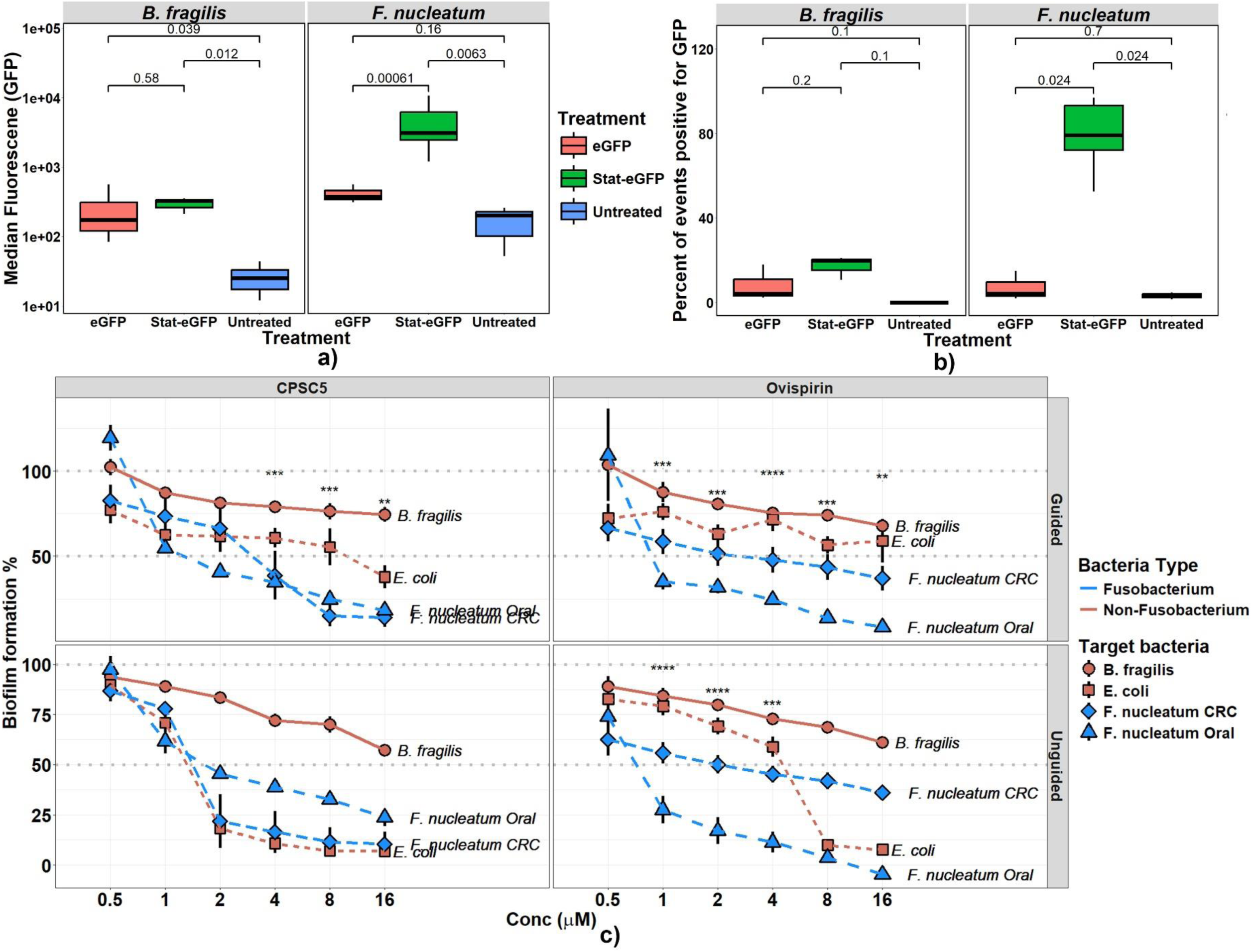
The Statherin derived guide peptide allows preferential attachment to *F. nucleatum* and exhibits targeted biofilm inhibition. (a) Median fluorescence intensity (MFI) of GFP in *B. fragilis* and *F. nucleatum* following treatment with eGFP, Stat-eGFP, and untreated controls. Statistically significant differences in GFP binding to *F. nucleatum* are indicated (p<0.05). Error bars represent the standard deviation of three independent experiments. (b) Percent of events positive for GFP, as measured by flow cytometry, comparing Stat-eGFP binding to *F. nucleatum* to *B. fragilis*. Statistical significance between treatments is noted (p<0.05). (c) Biofilm inhibition of *F. nucleatum* and *B. fragilis* strains in the presence of increasing concentrations of synthetic AMPs (Ovispirin and CPSC5) and their guided versions (Stat-Ovispirin and Stat-CPSC5). The inhibition is plotted as a percentage of biofilm formation. Error bars represent the mean ± standard deviation of three biological replicates. Statistical significance is indicated as *p*<0.05 (*), *p<0.01 (**), p<0.001 (*), and \**p*<0.0001 (**). Also see IC50 values calculated from the dose-response curve (Table 1)

**Table 1.** IC_50_ values of the synthetic AMPs – calculated by fitting biofilm inhibition assay data in a dose-response curve using *‘drc’* package (R)3.

| Bacteria |  | Ovispirin |  | CPSC5 |  |
| --- | --- | --- | --- | --- | --- |
|  |  | Unguided | Guided | Unguided | Guided |
| *Fusobacterium* | *F. nucleatum* (Oral) | 0.71 ± 1.71 µM | 0.91 µM | 2.18 ± 3.4 µM | 1.3 ± 4.36 µM |
|  | *F. nucleatum* (CRC) | 2.15 µM | 1.17 ± 1.04 µM | 1.42 ± 0.18 µM | 3.02 ± 0.69 µM |
| Non-*Fusobacterium* | *B. fragilis* | 23.54 ± 8.63 µM | >100 µM | 18.12 ± 6.37 µM | >100 µM |
|  | *E. coli* | 4.45 ± 0.35 µM | 17.26 ± 19.23 µM | 1.3 ± 0.1 µM | 9.68 ± 12.37 µM |

### Statherin-derived ligand guided AMP (gAMP) selectively inhibits biofilm formation of *F. nucleatum*

Next, using synthetic peptides (Ovispirin and CPSC5) and their guided analogs (stat-CPSC5 and stat-Ovispirin) we measured *in vitro* their biofilm inhibition capabilities. The synthetic peptides and their guided analogs were tested using 2 strains of *F. nucleatum*, one derived from CRC tumor samples (*F. nucleatum* SB-CTX3Tcol3) and another derived from an oral lesion (*F. nucleatum* subsp. *nucleatum* Knorr - ATCC 25586), along with well-described commensal strains of *B. fragilis* ATCC 43858 (anaerobic) and *E. coli* K12 ATCC 10798 (aerobic), as non-target bacteria candidates. Post incubation, using crystal violet assays for detection of biofilm formation, we show that both the AMPs and their corresponding gAMPs have similar patterns of inhibition against both the strains of *Fusobacterium* (**Fig 2C**), with their IC_50_ as low as 0.7 µM to as high as 3 µM (**Table 1**). While there was no change in MIC or IC_50_ observed in the gAMP compared to their AMPs against *F. nucleatum* (0.71-2.15 µM for Ovispirin, 0.91-2.3 µM for Stat-Ovispirin, *p=0.83*, 1.42-5.6 µM for CPSC5, 1.3-5.7 µM for Stat-CPSC5, *p=0.76*, ANOVA), when using the gAMP against the non-target bacteria, *B. fragilis* (23 µM for Ovispirin, >100 µM for Stat-Ovispirin, 18 µM for CPSC5, >100 µM for Stat-CPSC5, *p<0.01*, ANOVA) and *E. coli* (4.45 µM for Ovispirin, >30 µM for Stat-Ovispirin, 1.3 µM for CPSC5, >20 µM for Stat-CPSC5, *p<0.01*, ANOVA), we see a significant decrease in the inhibitory effect of the gAMPs compared to their AMPs, with the MIC and IC_50_ increasing significantly for the gAMPs against both the bacteria. This targeting effect was more pronounced against *B. fragilis,* with the gAMPs having greatly diminished inhibitory effect even beyond 32 µM and 64 µM of gAMP. For *E. coli*, the increase in IC_50_ was significant but not as pronounced as for *B. fragilis*, with a 4-fold to 7-fold increase in IC_50_. Thus, it can be concluded that the stat-CPSC5 and stat-Ovispirin were significantly more toxic towards *F. nucleatum* strains but not or less toxic compared to CPC5 and Ovispirin alone, or unguided. Against the non-target bacteria *B. fragilis* and *E. coli*, the gAMPs have significantly less toxicity compared to their AMPs when it comes to inhibition of biofilm formation. Overall, these data indicate that our gAMPs against FomA, have selective inhibition towards *F. nucleatum* over non-target bacteria. However, this differential toxicity towards the different bacteria species between the AMP and gAMP were not seen in cytotoxicity (MTT) assay against Caco2 and HT29 cells (**Fig S1**)

### Statherin-derived ligand-guided AMP (gAMP) differentially modulates growth kinetics of *F. nucleatum*

To better understand the effects of the gAMPs on growth of *F. nucleatum,* the growth kinetics of planktonic cultures of two non-*Fusobacterium* species – *E. coli* and *B. fragilis -* and two *Fusobacterium* strains - Oral and CRC - were incubated with both AMPs and their gAMP analogs. This analysis showed a pattern of differential inhibition (**Fig 3A-C**). For both the *Fusobacterium* strains, both the unguided and guided versions of the AMPs reduced the growth; starting between 2 and 8 µM, which continued for over 48 hrs and up to 72 hrs. In contrast, for the non-*Fusobacterium* species, only the unguided AMPs demonstrated any inhibitory effect below 8 µM, while the gAMPs either had no inhibition of growth or exhibited inhibition beyond 16-32 µM range. For *B. fragilis*, the bacteria that had the poorest response from all the AMPs, the guided CPSC5 and Ovispirin did not subdue growth, even at 32 µM, while the unguided version was effective at 8 µM. For *E. coli*, the AMPs started inhibiting growth even at 8 µM but the gAMPs only had inhibitory effects beyond 16 µM, with more pronounced inhibition at 32 µM. This analysis demonstrates that both AMPs and gAMPs inhibit the growth of *F. nucleatum* strains at low concentrations (<4 µM), making them clinically viable. Moreover, gAMPs show minimal toxicity towards non-*Fusobacterium* species compared to unguided AMPs, highlighting their selective toxicity and preferential action against *F. nucleatum* over the commensals.

**Fig 3.**
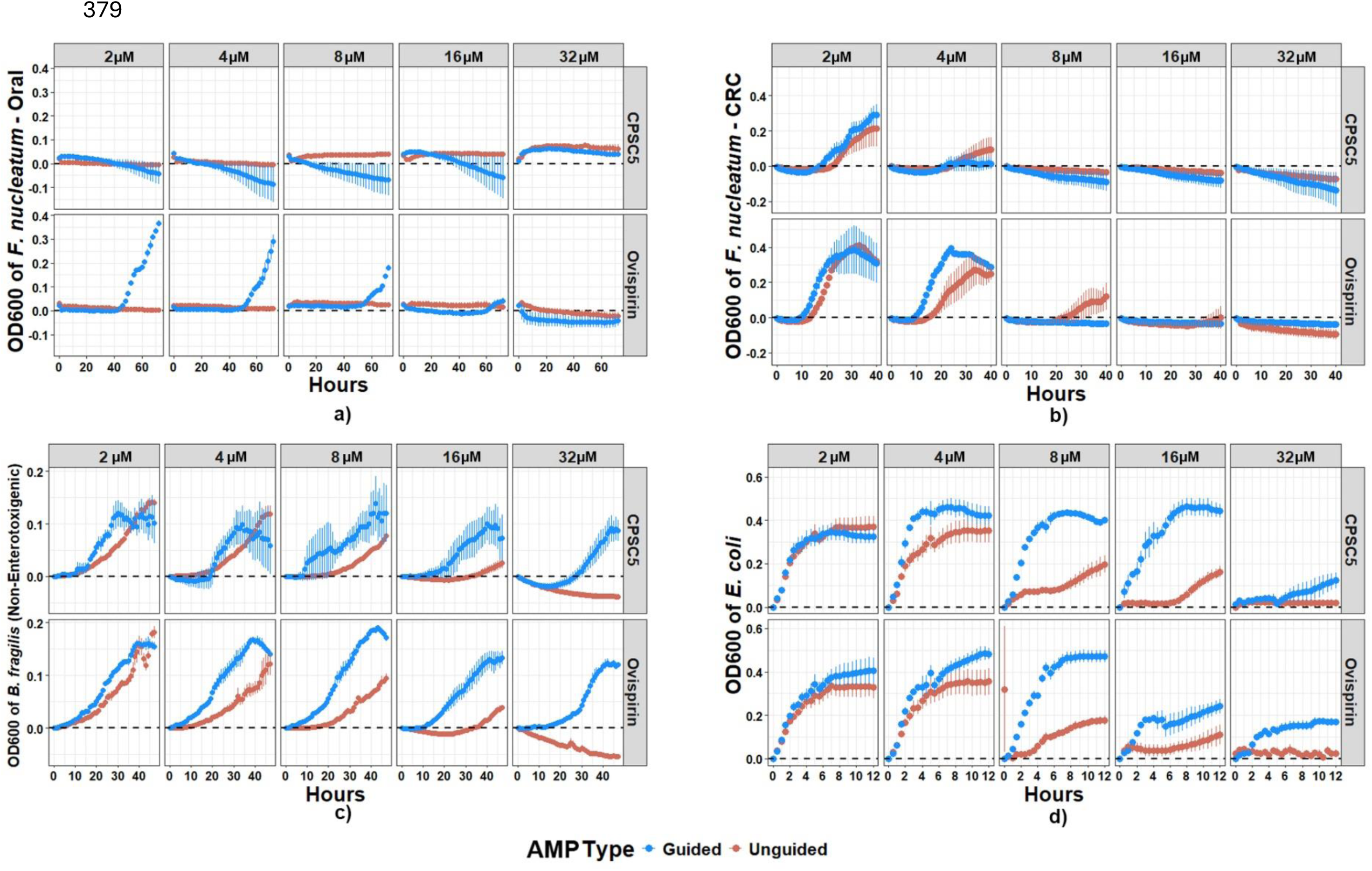
Selective inhibition of bacterial growth by guided antimicrobial peptides (gAMPs). Growth kinetics of planktonic cultures for two *Fusobacterium* strains—Oral/ATCC 25586 (a) and CRC/Strain 14 (b)—and two non-*Fusobacterium* species, *B. fragilis* (c) and *E. coli* (d), were measured in the presence of both unguided (red) and guided (blue) antimicrobial peptides (AMPs - CPSC5 and Ovispirin) across a concentration gradient (2 µM to 32 µM). Optical density (OD600) measurements were recorded over 60 hours (Oral strain) and 40 hours (CRC strain). Error bars represent standard deviation from three biological replicates.

### Cloning *Lactococcus lactis* with gAMP-Expressing Plasmids to Create Bioengineered Probiotics

Once the gAMPs demonstrated sufficient targeting abilities, the next step we took was creating the engineered probiotic strains expressing gAMPs. The open reading frames (ORFs) of the antimicrobial peptides (AMPs), CPSC5 and Ovispirin, along with their guided analogs (gAMPs), were cloned into the pMSP3545 plasmid (see Methods for details). This plasmid includes the nisin-inducible PNisA promoter and the usp45 signal peptide, which ensures the secretion of the peptides into the extracellular environment (**Fig. 4a**). The engineered plasmids were introduced into competent *Lactococcus lactis* MG1363 cells via electroporation. Successful cloning was confirmed through colony PCR using ORF-specific primers, followed by Sanger sequencing to verify the accuracy of the insertion. Based on these results, only the bioengineered probiotics expressing CPSC5 and Stat-CPSC5 were selected for further experiments, as the Ovispirin and stat-Ovispirin constructs did not yield consistent outcomes.

**Fig 4.**
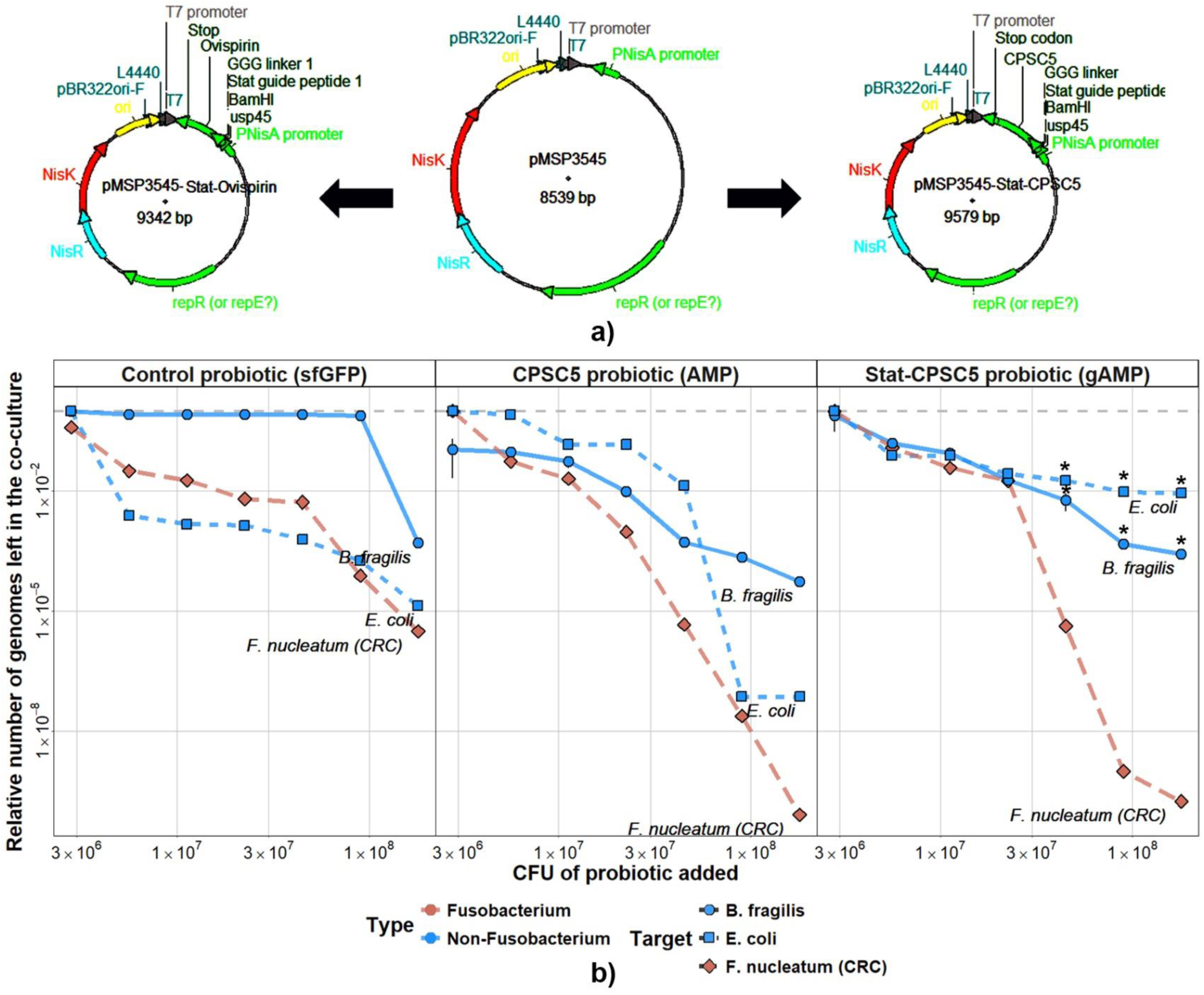
Engineered L. lactis with inducible plasmids demonstrates targeted inhibition of F. nucleatum compared to non-Fusobacterium species. (a) Schematic of the plasmids used to engineer L. lactis. The pMSP3545-derived plasmids contain the nisin-inducible PNisA promoter and the usp45 signal peptide for extracellular secretion of the peptides. The AMP/gAMP plasmids include ORFs for Ovispirin (left) or CPSC5 (right), with or without a Statherin-derived guide peptide. The control probiotic plasmid was constructed using the same backbone, replacing the AMP/gAMP ORFs with the superfolding GFP (sfGFP) ORF. Only CPSC5 and Stat-CPSC5 were successfully cloned into L. lactis. (b) Co-culture experiments with *F. nucleatum* (CRC strain), *Bacteroides fragilis*, and *E. coli* in the presence of control (sfGFP), AMP-expressing (CPSC5), or gAMP-expressing (Stat-CPSC5) *Lactococcus lactis* probiotics. The relative number of bacterial genomes remaining in the co-culture was quantified by species-specific qPCR after 24 hours of probiotic treatment, and plotted against the CFU of probiotic added. Error bars represent standard deviation from three biological replicates, and statistically significant differences (ANOVA, *p*<0.05) between guided and unguided AMPs are indicated by asterisks.

### Engineered *L. lactis* Expressing gAMP Exhibits Targeted Inhibition of *F. nucleatum* in Co-Culture Compared to Non-*Fusobacterium* Species

To further explore the specificity and potential clinical application of the bioengineered *L. lactis* probiotics, we conducted an *in vitro* co-culture experiment to assess the targeted inhibition of *F. nucleatum* compared to non-*Fusobacterium* commensal species. The bioengineered probiotics expressing CPSC5 (AMP) and Stat-CPSC5 (gAMP) demonstrated selective toxicity toward *F. nucleatum* when co-cultured with non-*Fusobacterium* commensal species. Both the AMP- and gAMP-expressing probiotics were co-cultured with a gradient of bacterial titers, ranging from 10^6 to 10^9 CFU/ml, of *F. nucleatum* (CRC strain), *B. fragilis*, and *E. coli*. After 24 hours of incubation, induced with 100 ng/ml of nisin in a 96-well plate format, the co-cultures were pelleted, and genomic DNA was extracted. Species-specific qPCR was used to quantify the target bacteria and estimate the remaining cell counts after incubation with the engineered probiotics. The results showed that the AMP-expressing probiotics inhibited the growth of all three bacterial species in a dose-dependent manner, with the most significant reduction observed in *F. nucleatum* (**Fig. 4b**). In contrast, the gAMP-expressing probiotics selectively inhibited *F. nucleatum* as the probiotic dose increased, while the inhibitory effects on *B. fragilis* and *E. coli* diminished significantly. The toxic effect of the gAMPs against *F. nucleatum* was 10^3 to 10^4 times greater than against *B. fragilis* and *E. coli* at higher probiotic titers (∼10^8 CFU/ml), a statistically significant difference (p < 0.05, ANOVA). Interestingly, some level of toxicity was observed across all three bacterial species, even in the control probiotic expressing super-folding GFP (sfGFP). This suggests that *L. lactis* may have inherent antimicrobial properties when co-cultured at high densities, possibly due to resource competition or innate toxicity, contributing to the observed inhibition.

### The gAMP Engineered Probiotic Maintains Microbial Diversity in a Polymicrobial Community

To determine whether the gAMP engineered probiotic could selectively target *F. nucleatum* while preserving the overall microbial diversity in a gut-like environment, we conducted a series of experiments using an *in vitro* human gut microbiome-derived polymicrobial community model. The efficacy of the bioengineered probiotics (AMP, gAMP, and Control) was tested within a polymicrobial community spiked with *F. nucleatum* (CRC strain) culture at 20% v/v in each well. This polymicrobial community was derived from anaerobic cultivation of rejuvenated fecal samples from three human subjects (FS04, FS16, and FS36), designed to mimic the composition of their gut microbiota. The communities were incubated with equal amounts of *F. nucleatum* and probiotic cultures—either AMP (CPSC5), gAMP (Stat-CPSC5), or Control (sfGFP). Samples were taken at 24 and 48-hour intervals, lysed and subjected to genomic DNA extraction using the phenol-chloroform method. The extracted gDNA was then analyzed using both qPCR with *F. nucleatum*-specific primers and 16S rRNA (V4) sequencing (Illumina) to assess the overall microbial composition. The 16S rRNA sequencing revealed that, while a diverse array of bacteria was present in the polymicrobial communities from each subject, the targeted modulation of *F. nucleatum* was most prominent in the samples treated with the engineered probiotics. This modulation was in stark contrast to the untreated controls, where *F. nucleatum* abundance increased considerably from around 10% to over 25% in 48 hrs (**Fig 5a & b**). Other prevalent genera, including *Enterococcus*, *Peptoniphilus*, *Eubacterium*, *Escherichia*, *Bacteroides*, *Sellimonas*, *Negati vicoccus*, and *Paraclostridium*, showed little to no change in relative abundance compared to *Fusobacterium*, indicating that the gAMP specifically modulates *F. nucleatum* without significantly affecting other members of the microbiota.

**Fig 5.**
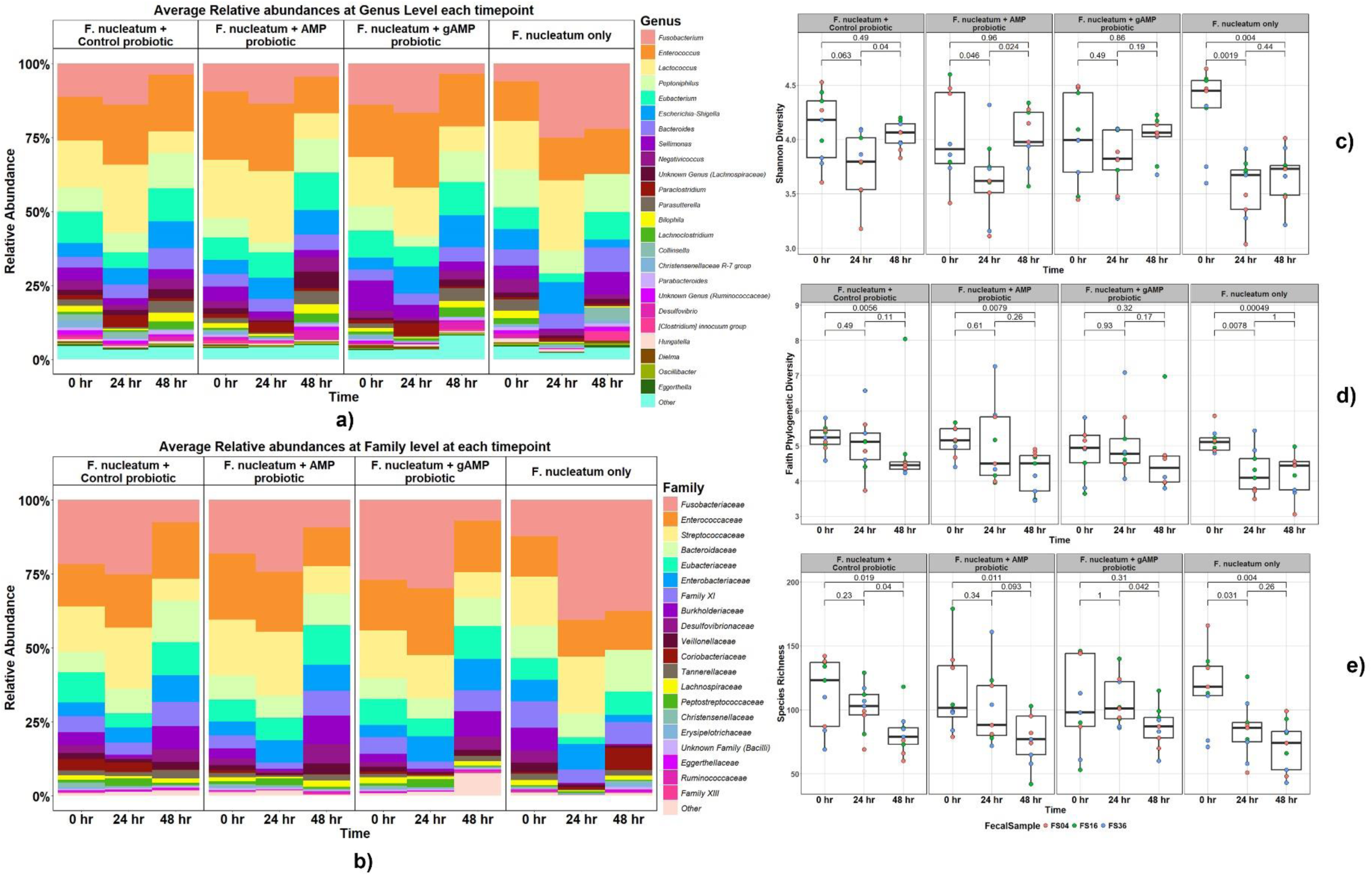
The gAMP probiotic maintained the diversity of the microbiota in the in vitro polmicrobial community. Heatmaps of the genus-level a) and family-level b) relative abundance. Polymicrobial community rejuvenated from human fecal microbiome sample spiked with *F. nucleatum* culture and then treated with bioengineered probiotics secreting CPSC5 (AMP), Stat-CPSC5 (gAMP) and sfGFP (Control) at a ratio of 6:1:1 respectively. b-d) Three alpha diversity indices – Shannon (a), Faith’s Phylogenetic diversity (b) and species richness (c), showing the ability of gAMP probiotic treatment to maintain the bioidiversity of the polymicrobial community post spiking with *F. nucleatum*, compared to AMP and control probiotics. No significant changes either between any consecutive timepoints or the overall timeframe. ANOVA was used for testing for significance between treatments with p<0.05.

Measuring the alpha diversity of the samples using three different indices revealed that all probiotic treatments preserved microbial community diversity after exposure to *F. nucleatum*. Among the treatments, the gAMP probiotic exhibited the smallest shift in microbial diversity from 0 to 48 hours. The Shannon index (**Fig. 5C**) indicates a decrease in diversity at 24 hours in all probiotic-treated samples, likely due to the proliferation of *F. nucleatum*. However, diversity began to recover in the following 24 hours. This decline at 24 hours was statistically significant in the control and AMP-treated samples but not in the gAMP-treated samples, with no significant differences in the recovery phase across all treatments. These results suggest that the gAMP probiotics caused the least disruption to microbial diversity, showing minimal and non-significant changes. In the samples that were spiked with *F. nucleatum* without probiotic treatment, diversity significantly declined at both 24 and 48 hours (p=0.0019 & 0.004 for Shannon Index, p=0.0078 & 0.0005 for Faith PD, p=0.031 & 0.004 for species richness, ANOVA). Interestingly, this dip and recovery pattern was not observed when analyzing the Faith Phylogenetic Diversity and Species Richness (**Fig. 5D-E**). Both of these indices showed a steady decline in diversity across all three probiotic treatments and in the *F. nucleatum*-only controls, with the gAMP-treated samples exhibiting the least drastic and non-significant changes in microbial diversity.

### The bioengineered probiotic effectively targets *F. nucleatum* in a gut microbiome-derived polymicrobial community without disrupting the overall microbiota

To assess whether the bioengineered gAMP probiotic could effectively reduce *F. nucleatum* in a complex gut microbiome-derived polymicrobial community without disrupting the overall microbial balance, we conducted a series of *in vitro* experiments. Using both 16S rRNA sequencing and qPCR, we found that the gAMP probiotic was highly effective in reducing *F. nucleatum* even within a complex polymicrobial community. Although all three probiotic variants demonstrated some antimicrobial activity, the gAMP probiotic produced the most significant reduction in *F. nucleatum* titers after 48 hours of incubation. Specifically, the *F. nucleatum* abundance, measured by Centered-Log Ratio (CLR) analysis, was lowest in the gAMP-treated samples compared to the AMP-treated ones, with a statistically significant reduction observed for the gAMP probiotic (**Fig. 6A**) (p=0.031, ANOVA). In the probiotic-treated samples, there was a slight increase in *F. nucleatum* CLR at the 24-hour mark, followed by a notable decline, which was significant only in the gAMP-treated samples. In contrast, samples treated only with *F. nucleatum* exhibited a steady increase in CLR over time. Interestingly, these significant reductions in *F. nucleatum* abundance were not fully reflected in the qPCR results, where similar reductions were observed across all probiotic treatments. Only the untreated control samples showed a significant increase in *F. nucleatum* titer at both time points (p=4.1e-05, ANOVA) (**Fig. 6A**). This discrepancy between the two methods may be attributed to the nature of the measurements: qPCR focuses solely on the number of gene copies for the target species, whereas CLR analysis accounts for the relative abundance of *F. nucleatum* in relation to all other bacteria in the community, offering a more comprehensive view of its presence within a polymicrobial environment.

**Fig 6.**
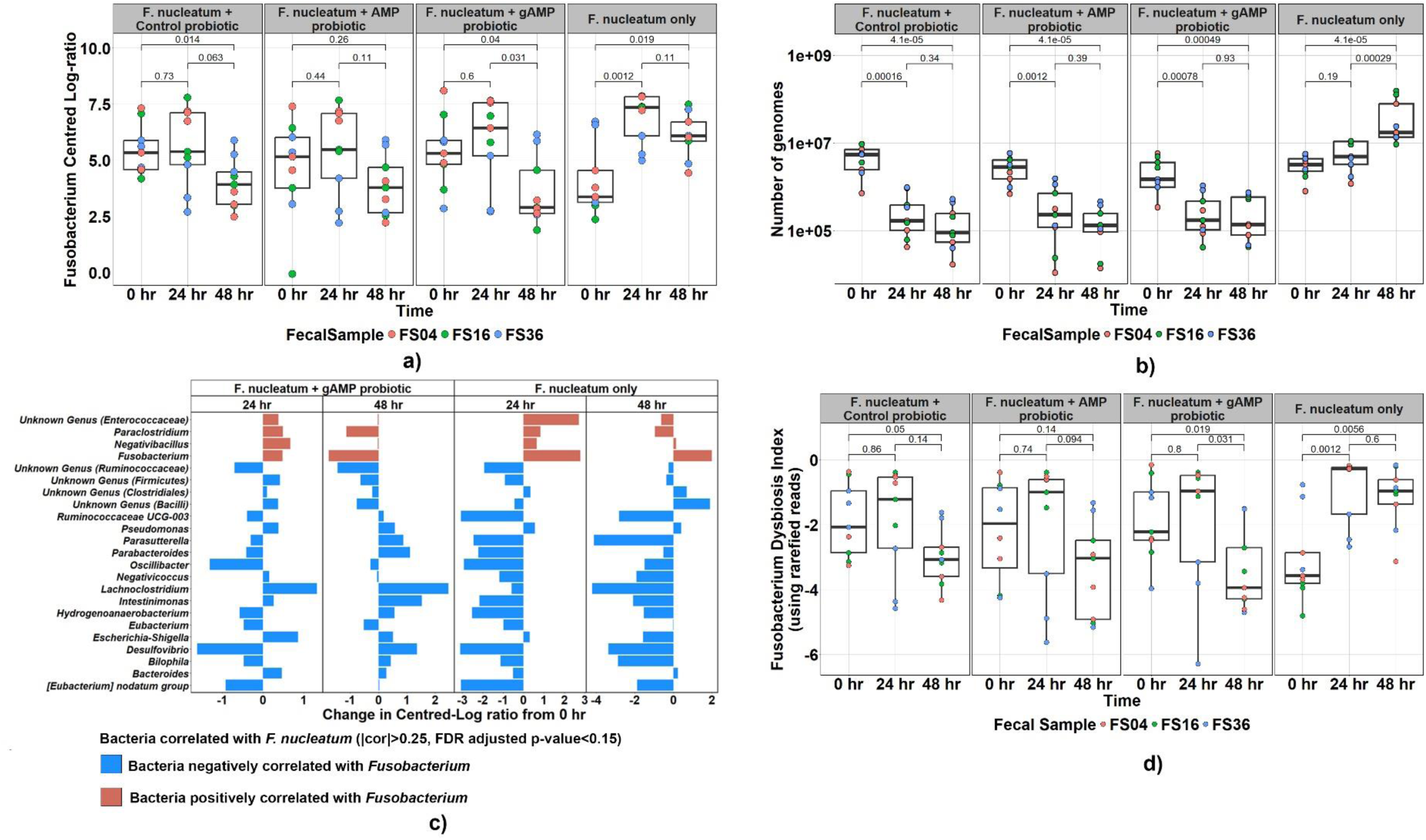
The bioengineered probiotic effectively targets *F. nucleatum* in a gut microbiome-derived polymicrobial community without disturbing the overall microbiota. *(a)* The effect control probiotics, AMP, or gAMP probiotic on reduction in *F. nucleatum* titers after 48 hours in an in vitro polymicrobial community. The abundance of *F. nucleatum* was measured by Centered-Log Ratio (CLR), (ANOVA).b) *F. nucleatum* titer measure by qPCR of the sample samples. The y-axis gives the approximate number of *F. nucleatum* left in the sample at that timepoint after treatment with the given probiotic. (ANOVA, p<0.05) *(c)* Compositionality Corrected by REnormalization and PErmutation (CCREPE) analysis of bacterial genera that were positively (red) or negatively (blue) correlated with *F. nucleatum* abundance. This analysis was used to create the Fusobacterium Dysbiosis Index. *(d)* The effect of probiotic, AMP, or gAMP probiotics on community structure, as indicated by the Fusobacterium Dysbiosis Index, which was calculated by comparing the abundance of *F. nucleatum* and its positively correlated genera against the negatively correlated genera.(Dysbiosis Index = |correlation| > 0.25, FDR-adjusted p < 0.15)

A Compositionality Corrected by REnormalization and PErmutation (CCREPE) analysis was performed to identify bacterial species either positively or negatively correlated with *F. nucleatum* abundance (|correlation| > 0.25, FDR-adjusted p < 0.15) across the samples. This allowed us to determine which bacteria increased as *F. nucleatum* levels rose, and which decreased in response. In the gAMP probiotic-treated samples, bacteria positively correlated with *F. nucleatum* showed a consistent reduction, while those negatively correlated were either maintained or increased in abundance at both time points (**Fig. 6C**). In contrast, the samples treated only with *F. nucleatum* exhibited the opposite trend. This information was used to develop the Fusobacterium Dysbiosis Index (**Formula 1**), which quantifies the extent of dysbiosis caused by introducing *F. nucleatum* into the community. Using this index, the gAMP probiotics showed the most effective performance in vitro (**Fig. 6D**). The gAMP-treated samples exhibited a significant reduction in dysbiosis, indicating that not only did the gAMP probiotics reduce *F. nucleatum* abundance, but they also preserved microbial diversity more effectively than AMP probiotics by selectively targeting the pathogen.

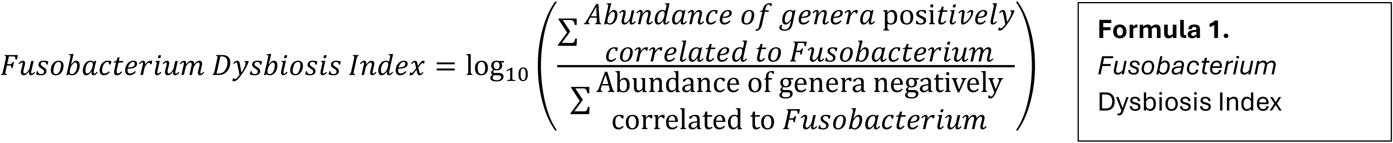

Although all three probiotic treatments showed a similar pattern of a slight increase in dysbiosis at 24 hours followed by a reduction at 48 hours, the decrease in the gAMP-treated samples was significantly lower than at both the 0-hour and 24-hour time points—an effect not seen with the AMP or control probiotics. These results confirm that the gAMP probiotic expressing Stat-CPSC5 demonstrates preferential antimicrobial activity against *F. nucleatum*, consistent with findings from both monoculture and polymicrobial community experiments.

## Discussion

Our research presents critical insights into the specificity and efficacy of Statherin-derived guide peptides as a first step in creating an engineered probiotic selectively targeting *F. nucleatum* highlighting the potential for targeted antimicrobial therapies. Our study investigates the implications of these findings, particularly in relation to the selective bacterial targeting, inhibition of biofilm formation, modulation of bacterial growth, and the preservation of microbial diversity within complex polymicrobial communities.

By utilizing a statherin-derived guide peptide fused to an enhanced green fluorescent protein (eGFP), we observed preferential attachment to *F. nucleatum* over *B. fragilis*, a non-Fusobacterium commensal anaerobe [49]. Flow cytometry revealed significantly higher fluorescence in *F. nucleatum* treated with stat-eGFP compared to both native eGFP and untreated controls, indicating a specific interaction [50]. In contrast, *B. fragilis* exhibited no significant difference between native eGFP and stat-eGFP treatments, confirming the specificity of the statherin-derived peptide in guiding attached molecules to bind selectively to *F. nucleatum*. This specificity is mediated by the interaction between the Statherin-derived peptide and the membrane protein FomA, as shown by previous studies in which *F. nucleatum* mutants lacking FomA lost binding specificity to the guide peptide ligand [39]. This specificity is crucial in designing targeted therapies that minimize the impact on non-target microbiota, thereby reducing potential off-target effects and preserving the delicate balance within the gut microbiome [51, 52].

Biofilms provide a protective environment for bacteria, increasing their resistance to antimicrobial agents and immune responses [53]. Our biofilm inhibition assays further demonstrated that statherin-derived ligand-guided antimicrobial peptides (gAMPs), including stat-CPSC5 and stat-Ovispirin, retained strong inhibitory effects against *F. nucleatum* biofilms while showing reduced efficacy against non-target bacteria such as *B. fragilis* and *E. coli*. This selective inhibition is particularly important in the context of treating bacterial infections linked to biofilm formation. The ability of gAMPs to selectively inhibit *F. nucleatum* biofilms without affecting commensal bacteria suggests their potential as therapeutic agents for biofilm-associated infections, particularly in the oral cavity and gastrointestinal tract [54, 55].

Growth kinetics data also supported the specificity of gAMPs, showing that both guided and unguided AMPs inhibited *F. nucleatum* at low clinically relevant concentrations. However, gAMPs exhibited significantly lower toxicity toward non-*Fusobacterium* bacteria. For example, in *B. fragilis*, gAMPs had a negligible inhibitory effect even at higher concentrations, whereas unguided AMPs were effective at much lower doses. This selective growth inhibition emphasizes the therapeutic potential of gAMPs, where it is essential to target pathogenic bacteria without disrupting the broader microbial community [51, 52]. Such precision could prevent the adverse effects commonly associated with broad-spectrum antibiotics, such as dysbiosis and the emergence of antibiotic-resistant strains. These results also confirm that the gAMP strategy is effective against both planktonic and biofilm forms of the target pathogen.

The creation of bioengineered probiotics expressing gAMPs introduces an innovative approach to delivering targeted antimicrobial therapy within the gut. There has been recent research done using the approach of targeted antimicrobial peptide against *F. nucleatum* using the FomA protein [56], but it did not offer a suitable delivery platform for delivery of the peptides in situ of infection. Engineered *Lactococcus lactis* developed by us, expressing stat-CPSC5, exhibited selective toxicity towards *F. nucleatum* when co-cultured with non-target bacteria. The significant reduction in *F. nucleatum* populations, as indicated by quantitative PCR and species-specific genome quantification, was observed without comparable effects on *B. fragilis* and *E. coli*. This differential targeting is advantageous for maintaining a healthy gut microbiota while combating pathogens. Additionally, the inherent antimicrobial activity of *L. lactis* suggests it could serve as an effective vehicle for delivering therapeutic agents, enhancing its utility in microbial therapy.

One of the most significant findings of this study is the ability of gAMP-expressing probiotics to preserve the diversity of a polymicrobial community while effectively reducing *F. nucleatum* levels. 16S rRNA sequencing showed that gAMP probiotics maintained microbial diversity to a greater extent than AMP or control probiotics. Alpha diversity (Shannon Index) demonstrated that while all samples showed a decrease in diversity at 24 hours, likely due to *F. nucleatum* proliferation, the gAMP probiotic-treated samples exhibited a less significant decrease and a more complete recovery by 48 hours. This suggests that gAMPs can selectively reduce pathogenic bacteria while minimizing collateral damage to the overall microbial community.

Moreover, compositional analysis using CCREPE [48] revealed that gAMP probiotics were most effective in reducing the dysbiosis index, a measure of microbial imbalance caused by *F. nucleatum* overgrowth [57]. This reduction in dysbiosis correlated with the selective decrease in bacteria positively associated with *F. nucleatum* and the preservation or increase of bacteria negatively associated with *F. nucleatum*. These findings are significant because maintaining microbial diversity and function is essential for gut health, and significant and/or prolonged disruptions can lead to conditions such as inflammatory bowel disease, cancer and other metabolic disorders [58].

Our findings have broad implications for developing targeted antimicrobial therapies. The specificity of Statherin-derived guide peptides for *F. nucleatum* opens up the possibility of using gAMPs in clinical settings to treat infections associated with this pathogen, particularly in the context of colorectal cancer, periodontal disease, and even reproductive diseases such as pre-term birth and endometriosis [14, 59–62]. Additionally, the use of bioengineered probiotics as a delivery system offers a promising avenue for administering these therapies in a controlled and sustained manner. Future research should focus on translating this therapeutic approach to in vivo models, where bioengineered *L. lactis* synthesizing and secreting gAMP could be administered orally for gut colonization, allowing the peptides to escape gastric proteolysis and exert antimicrobial effects in situ by preferentially binding to *F. nucleatum*’s FomA surface protein via the Statherin-derived guide peptide. (**Fig 7**).

**Fig 7.**
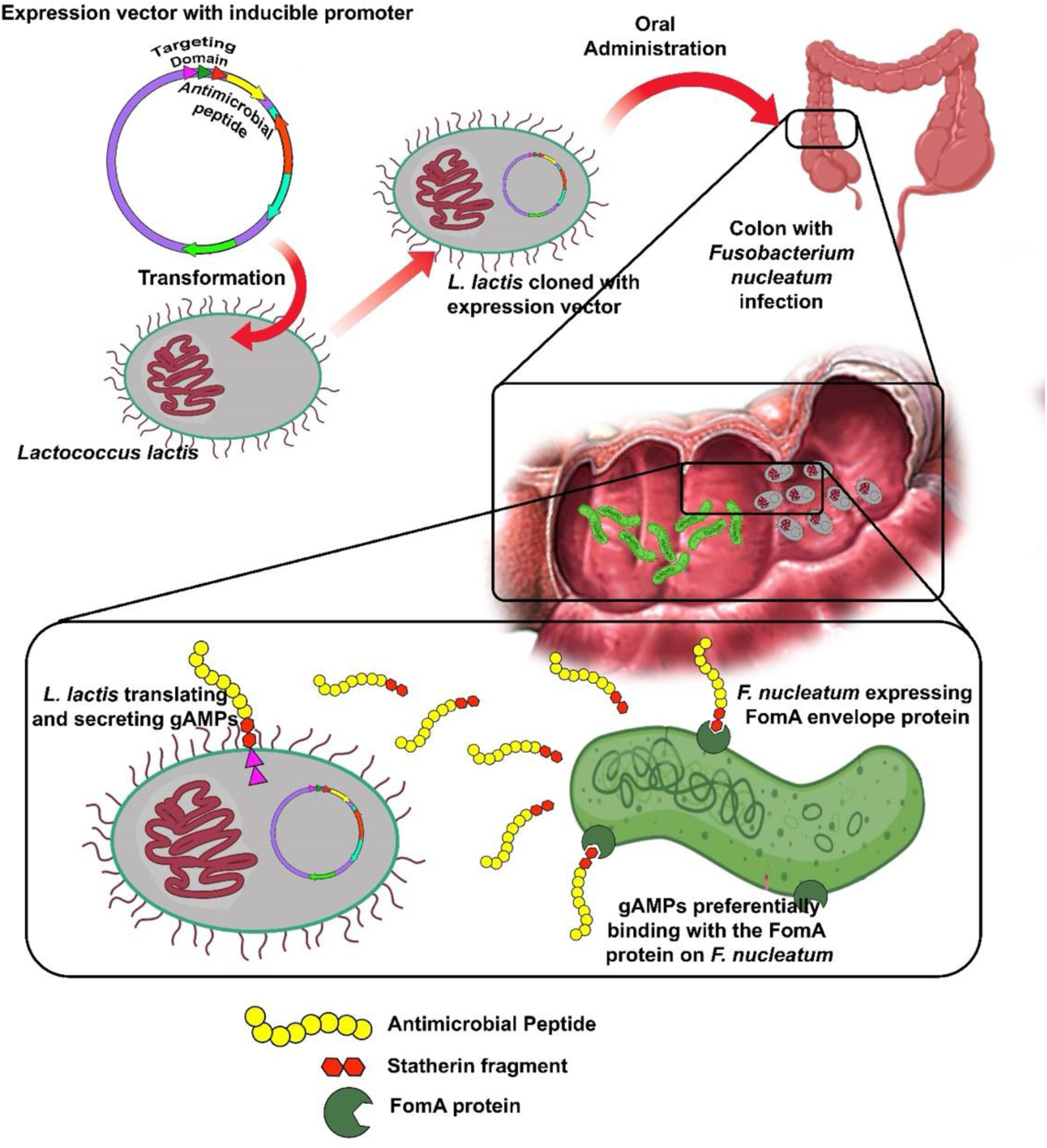
Model for in vivo application of bioengineered probiotics as a therapeutic. The probiotic schematic representation illustrates the engineered probiotic mechanism. The probiotic Lactococcus lactis is bioengineered with a Nisin-inducible plasmid that includes the PNisA promoter and the usp45 secretion signal peptide. This vector enables the expression and secretion of antimicrobial peptides (AMPs) or guided antimicrobial peptides (gAMPs) in the gut upon oral administration. The statherin-derived guide peptide on the gAMPs targets the FomA membrane protein expressed by F. nucleatum. Once secreted, the gAMPs preferentially bind to FomA on F. nucleatum, allowing targeted antimicrobial activity and thereby reducing F. nucleatum colonization in the infected colon without affecting non-target bacteria.

Furthermore, the ability to maintain microbial diversity while selectively targeting pathogens could revolutionize the treatment of bacterial infections. As indicated, the current antibiotic therapies often disrupt the gut microbiota, leading to adverse effects and promoting the development of antibiotic resistance. The use of gAMPs, particularly when delivered via probiotics, could mitigate these issues by providing a more targeted approach that preserves the beneficial components of the microbiota in an easily delivered oral therapeutic.

In conclusion, this study demonstrates the effectiveness of Statherin-derived guide peptides in selectively targeting *F. nucleatum*, inhibiting its biofilm formation, and modulating its growth kinetics without significantly affecting non-target commensal bacteria. The bioengineered probiotics expressing gAMPs further showed promise in reducing *F. nucleatum* within a complex polymicrobial community while maintaining microbial diversity. These findings lay a strong foundation for the development of targeted antimicrobial therapies that could offer a more precise and less disruptive alternative to traditional antibiotics.

## Data Availability

The 16s rRNA amplicon sequences of the polymicrobial community derived from the fecal samples used n the study are available in NCBI SRA database under the BioProject ID PRJNA1179861.

## Acknowledgements

We acknowledge the help of members of the laboratory of Dr. Christopher M. Kearney, Department of Biology, Baylor University for assistance with the development of the guided antimicrobial peptide and bioengineered probiotic technology including Dr. Mikaeel Young, Dr. Patrick Ortiz, and Dr. Toslim Mahmud for help in procuring bacteria strains and performing assays. We acknowledge the help of Dr. Emma Allen-Vercoe, Department of Molecular and Cellular Biology, University of Guelph, ON, Canada and Dr. Susan Bullman, Department of Immunology, MD Anderson Cancer Center, Texas, USA for sharing their clinically sourced strains of *Fusobacterium nucleatum* with us.

---

**Fig S1.**
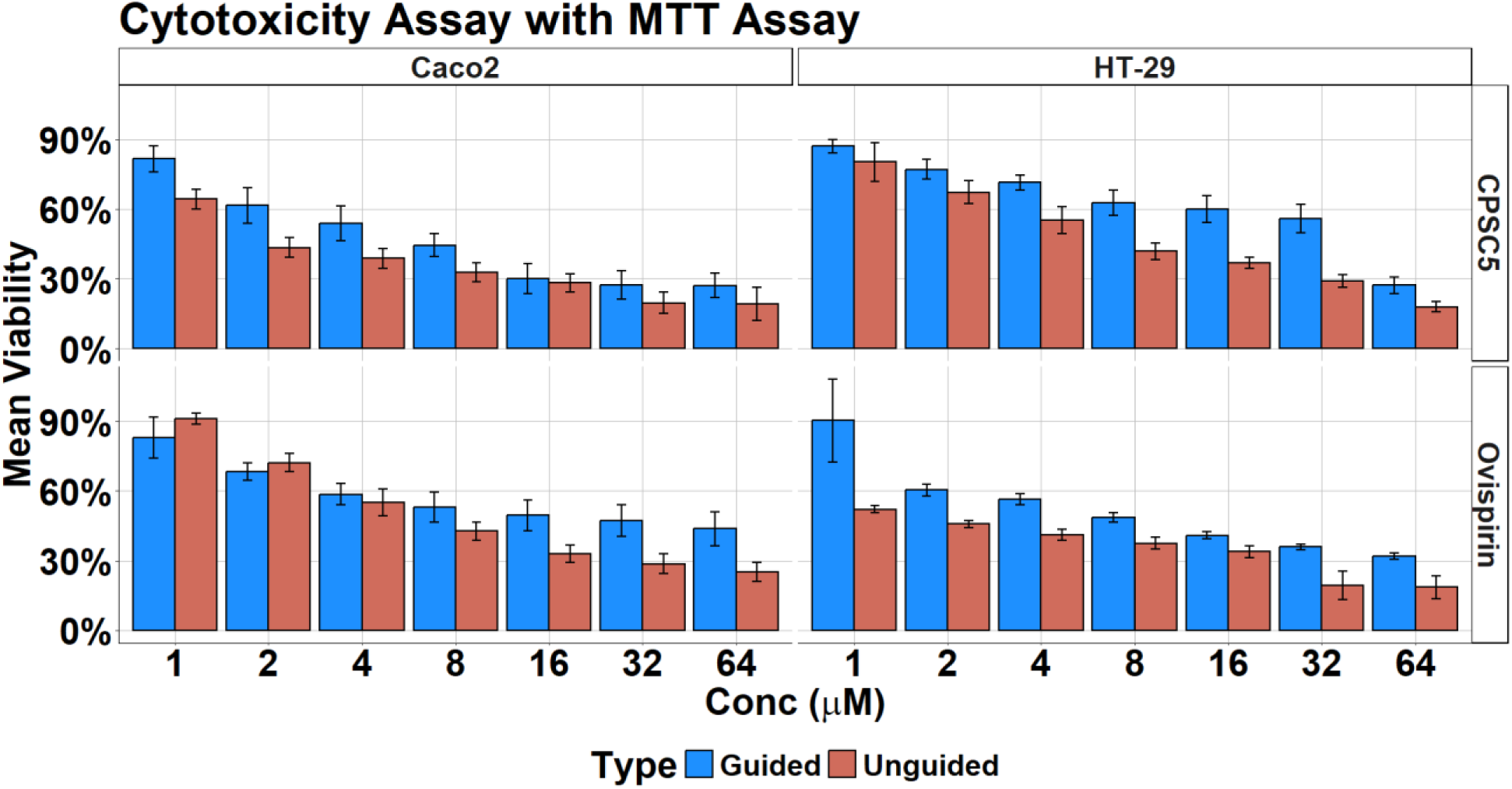
MTT Assay of Ovispirin and CPSC5 and their Stat-guided analogs against Caco2 and HT29 cells

**Table S1.**
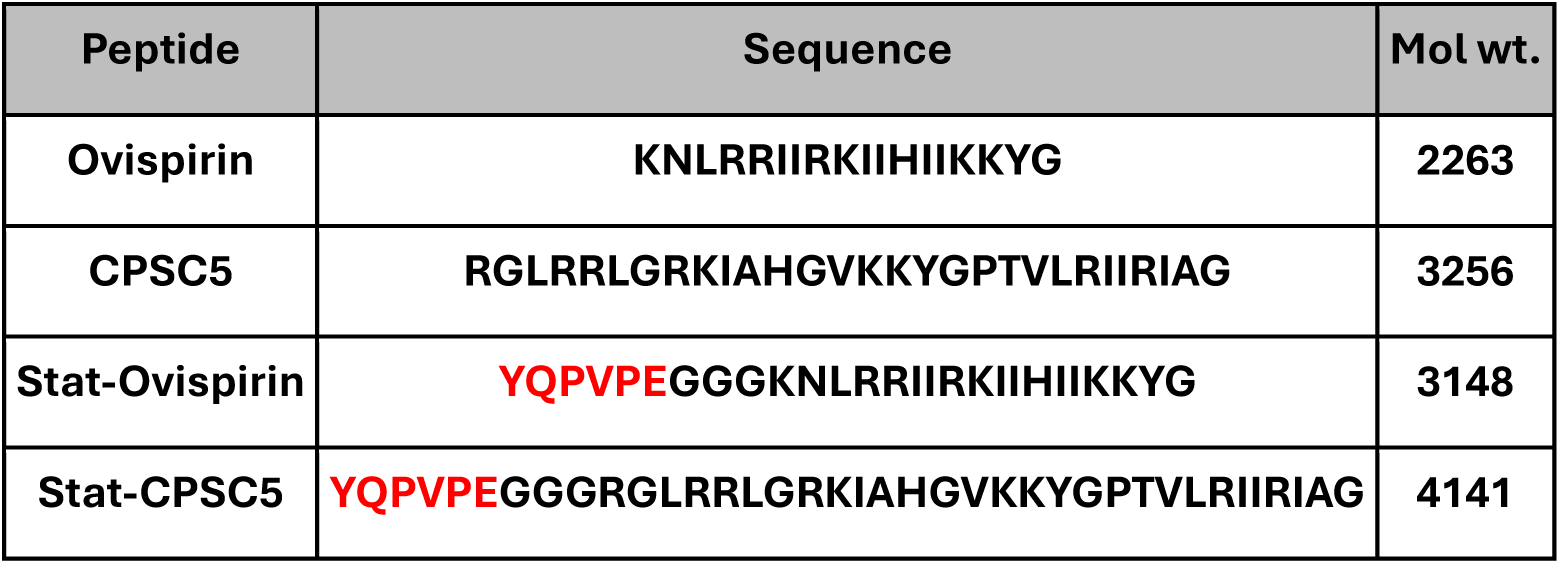
AMPs used in the study – Ovisiprin and Cathelin-derived Peptide SC5. The statherin derived guide peptide (red) is attached to the N-terminus of the AMPs, separated by a –GGG-linker, to create the corresponding guided AMPs (gAMPs).

